# Antiviral immunity within neural stem cells distinguishes viral strain differences in forebrain organoids

**DOI:** 10.1101/2024.07.09.602767

**Authors:** Christine Vazquez, Seble G. Negatu, Carl D. Bannerman, Sowmya Sriram, Guo-Li Ming, Kellie A. Jurado

## Abstract

Neural stem cells have intact innate immune responses that protect them from virus infection and cell death. Yet, viruses can antagonize such responses to establish neuropathogenesis. Using a forebrain organoid model system at two developmental time points, we identified that neural stem cells, in particular radial glia, are basally primed to respond to virus infection by upregulating several antiviral interferon-stimulated genes. Infection of these organoids with a neuropathogenic Enterovirus-D68 strain, demonstrated the ability of this virus to impede immune activation by blocking interferon responses. Together, our data highlight immune gene signatures present in different types of neural stem cells and differential viral capacity to block neural-specific immune induction.

## Introduction

Enterovirus-D68 (EV-D68) is a re-emerging, single-stranded, positive-sense RNA virus that was first isolated from four children with respiratory disease in the 1960s [1]. Intriguingly, since 2014, EV-D68 has been associated with a polio-like paralysis called acute flaccid myelitis (AFM) as biennial AFM cases have coincided with EV-D68 outbreaks in the United States [2]. Epidemiological surveillance and virus genome sequencing have confirmed the divergence of EV-D68 strains from Fermon, a prototypic strain from the 1960s, suggesting virus evolution and differential strain tropism [3]. Studies to determine the capacity of contemporary and prototypic EV-D68 strains to infect cells of the central nervous system (CNS) are conflicting [4, 5]. Given the emergence of respiratory viruses associated with neurological disease, understanding the neural tropism and viral factors associated with pathogenesis is important.

Brown and colleagues showed that strain VR1197, representative of a prehistoric EV-D68 strain similar to the prototypic strain Fermon, replicated at low levels in differentiated neuroblastoma cells and primary human fetal brain-derived neurons [4]. However, several other studies have suggested that EV-D68 neurotropism is not a recently acquired phenotype as contemporary circulating EV-D68 strains and two 1962 strains (Fermon and Rhyne) replicate in various murine and human neuronal systems [5, 6]. In addition to the inconclusive host cellular tropism of EV-D68, the viral receptor remains under investigation. Thus, understanding how the different EV-D68 strains interact within the cellular environment to dictate virus strain differences is important towards our goal of understanding the role immune responses play in protecting from neurological disease.

Throughout CNS development, the human brain is largely protected from virus infection through antiviral innate immune signaling and protective barriers [7]. Given the relative post-mitotic nature of neurons and other cells of the CNS, immune signaling serves a vital role in protecting these limited cells from excessive inflammation and cellular death. Early in development, stem cells serve as gatekeepers, guarding the developing brain. However, neurotropic viruses may bypass such defenses to enter the CNS and cause neurological disease [8, 9]. CNS-specific immunity and viral immune evasion mechanisms are only beginning to be uncovered.

We hypothesized that type I antiviral signaling within cells of the CNS could contribute to EV-D68 strain differences. To model the heterogeneous complexity of the CNS, we utilized a recently established forebrain organoid model at two developmental time points and identified immune regulation within different neural stem cell subtypes by different EV-D68 strains. We found that neural stem cells basally express higher levels of several interferon stimulated genes (ISG); neuropathogenic EV-D68 can block upregulated *IFITM1* defenses; and that different neural lineages contain, diverse immune gene signatures, which may protect against virus infection.

## Results

### SH-SY5Y cells are permissive to neuropathogenic and non-neuropathogenic EV-D68 strains

Several studies have sequenced circulating EV-D68 strains in many countries, identifying virus mutations that have diverged from the historic Fermon strain (illustrated in **Figure 1A**). However, whether EV-D68 mutations have conferred CNS permissiveness remains unclear. Given the inconsistent replication kinetics reported in the literature, we sought to determine whether Fermon and a contemporary neurotropic EV-D68 strain, MO/14-18947 (defined throughout as MO), could infect either undifferentiated or differentiated SH-SY5Y cells. SH-SY5Y cells have historically been used to investigate EV-D68 replication dynamics [4]. To differentiate SH-SY5Y cells, we utilized a previously established 18-day protocol dependent upon addition of retinoic acid and gradual depletion of fetal bovine serum [10]. Temperature is known to influence EV-D68 replication kinetics with 33°C, that of the upper respiratory tract, being more suitable; however, there is still infectious virus production at 37°C [11, 12]. Our investigation sought to examine the mechanisms underlying EV-D68 replication and immune regulation within the CNS, where the human temperature is 37°C. Thus, we performed our infections at 37°C. To determine virus permissiveness, we first infected undifferentiated SH-SY5Y cells at varying multiplicities of infection (MOI) and measured viral VP1 protein expression via immunofluorescence (**Figure 1B)**, viral RNA copies (**Figure 1C),** and viral titer **(Figure 1D).** We found that both Fermon and MO could infect and replicate in undifferentiated SH-SY5Y cells, even at the lowest MOI. We next differentiated SH-SY5Y cells and infected them with Fermon and MO at varying MOIs. Similar to the undifferentiated cells, we found that both Fermon and MO expressed viral VP1 protein (**Figure 1E)**, replicated to detectable RNA copies **(Figure 1F)**, and produced infectious virus titer **(Figure 1G).** Together, these data support the notion that SH-SY5Y cells are not ideal models to recapitulate the EV-D68 strain differences in the CNS as non-neuropathogenic Fermon was capable of infecting and replicating within them.

**Figure 1.**
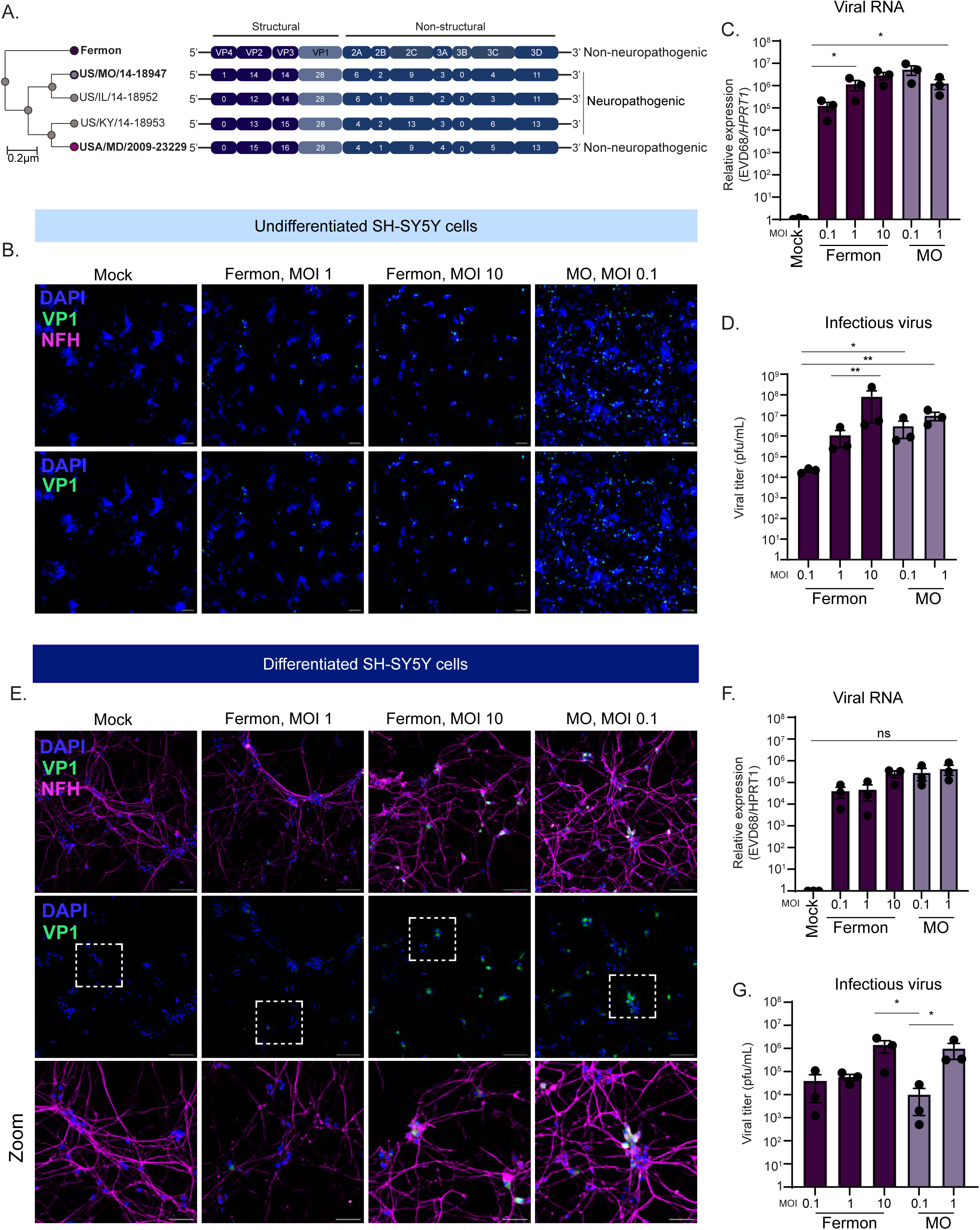
Prototypic Fermon strain replicates in SH-SY5Y cells. **(A).** Alignment and phylogenetic tree of several EV-D68 non-neuropathogenic and neuropathogenic strains, with the prototypic Fermon strain serving as reference strain. This schematic was adapted from [11]. Phylogenetic tree from viral genome sequences was done using the NCBI phylogenetic tree software, and the protein sequences were aligned using Clustal Omega. **(B).** Immunofluorescence images of undifferentiated SH-SY5Y cells mock-infected or infected with Fermon or MO viruses at the indicated multiplicity of infection (MOI) at 24 hours post infection (hpi). Cells were stained for DAPI (nuclei, blue), VP1 (green), and neurofilament-H (NFH, magenta). Scale bar = 100 µm. **(C).** RT-qPCR analysis of RNA harvested from undifferentiated SH-SY5Y cells mock-infected or infected with Fermon or MO at the indicated MOIs for 24 hours. The data are normalized to mock-infected cells and are presented as relative expression of EV-D68 to *HPRT1* with Mock set to 1. Data are presented as means ± SEM (n = three biological replicates). **(D).** Plaque-forming assay of supernatants from undifferentiated SH-SY5Y cells infected with either Fermon or MO at the indicated MOIs for 24 hours. Data are presented as means ± SEM (n = three biological replicates). N.D. = not detected **(E).** Immunofluorescence images of differentiated SH-SY5Y cells mock-infected or infected with Fermon or MO viruses at the indicated multiplicity of infection (MOI) at 24 hpi. Cells were stained for DAPI (nuclei, blue), VP1 (green), and neurofilament-H (NFH, magenta). Scale bar = 100 µm. **(F).** RT-qPCR analysis of RNA harvested from differentiated SH-SY5Y cells mock-infected or infected with Fermon or MOI at the indicated MOIs for 24 hours. The data are normalized to mock-infected cells and are presented as relative expression of EV-D68 to *HPRT1* with Mock set to 1. Data are presented as means ± SEM (n = three biological replicates). N.D. = not detected **(G).** Plaque-forming assay of supernatants from differentiated SH-SY5Y cells infected with either Fermon or MO at the indicated MOIs for 24 hours. Data are presented as means ± SEM (n = three biological replicates). Statistical analysis performed with one-way analysis of variance (ANOVA).

### Developmentally less mature forebrain organoids are selectively permissive to neuropathogenic EV-D68

To attempt to better model EV-D68 strain differences observed in the human population *in vitro,* we turned to the recently established forebrain organoid model system. Forebrain organoids display the complex cellular environment observed in the developing human cortex, with sustained maintenance in culture permitting development of more mature neuronal subtypes. [13] **(Figure 2A)**. To model a heterogeneous cell population and emergence of maturing neuronal cells, we cultured forebrain organoids until either day *in vitro* (DIV) 35 (early) or 85 (late) from the human induced pluripotent stem cells [14–16]. Early organoids are characterized by the appearance of cellular niches of stem and progenitor cells, or neural rosettes, that line the periphery of the organoid **(Figure 2B)**. On the other hand, late organoids contain few neural rosette structures **(Figure 2B)**. Both early and late organoids were mock, Fermon-, or MO-infected for 24 hours. At 24 hours post-infection (hpi), infection inoculum was removed, and media was changed daily until 7 days post infection (dpi), at which time both organoids and culture supernatant were harvested for analysis. We observed viral RNA copies **(Figure 2C)** in both Fermon- and MO-infected organoids, but infectious viral titer **(Figure 2D)** was limited to MO-infected organoids only. At the late time point, however, we found that both Fermon and MO are capable of replicating **(Figure 2E)** and producing infectious titer **(Figure 2F).** We next examined viral VP1 protein via immunofluorescence of early and late organoids and saw that only MO-infected early organoids contained VP1-positive cells **(Figure 2G)** whereas both Fermon and MO-infected late organoids expressed viral antigen **(Figure 2H)**. To address the concern of reproducibility amongst various human iPSCs, we validated findings with forebrain organoids generated from another iPSC line, WTC11 [17], and found similar phenotypes **(Supplemental Figure 1)**. These data establish forebrain organoids as an *in vitro* system that phenocopies the neuronal replication kinetics observed in the human population.

**Figure 2.**
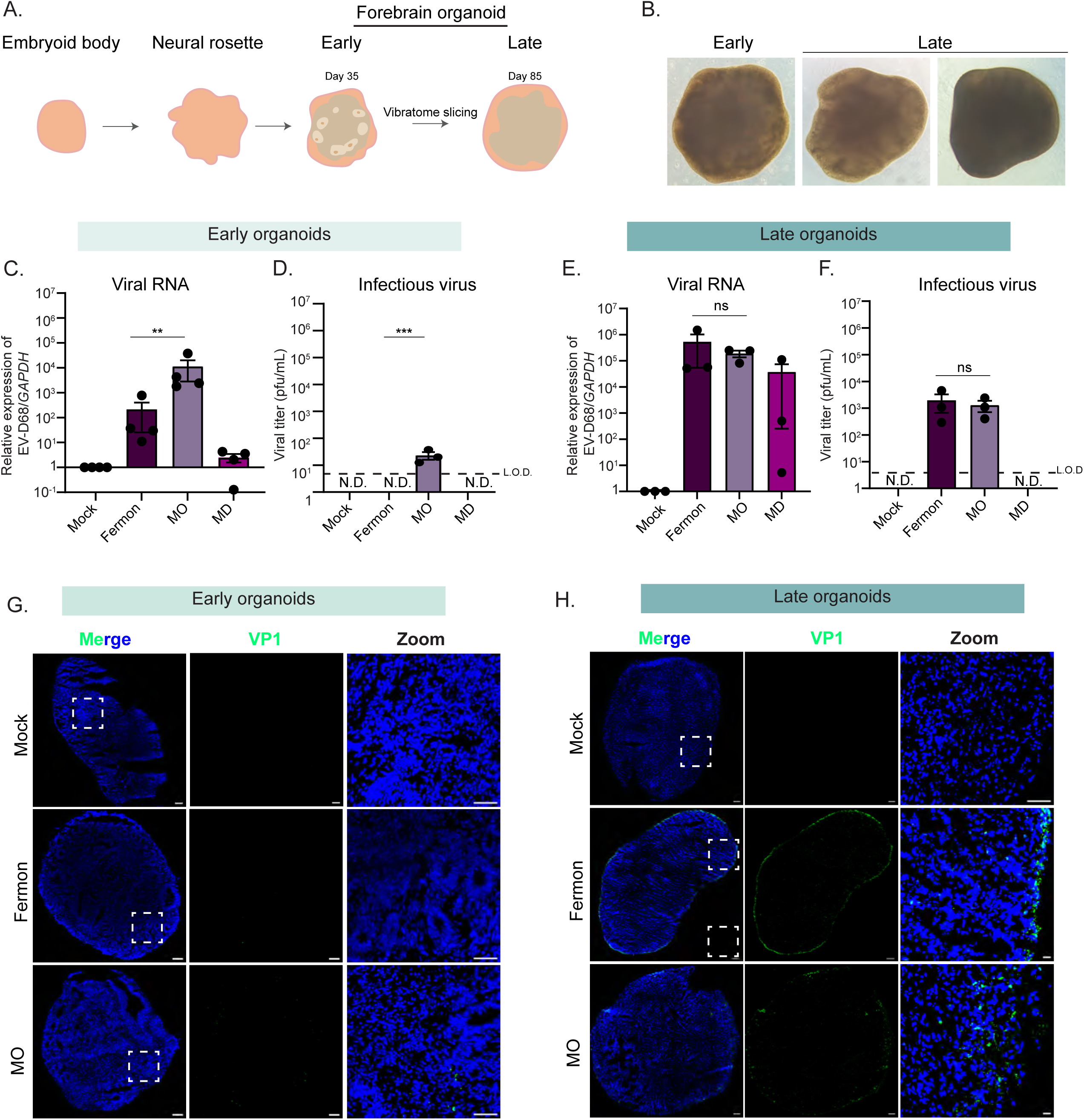
Differential Enterovirus-D68 replication in early and late forebrain organoids. **(A).** Schematic of organoid structures. **(B).** Bright field images taken at 40X from organoids at day *in vitro* (DIV) *35* (early) and DIV85 (late) **(C-H).** Early and late organoids were infected mock-infected or infected with Fermon, MO, and MD viruses for 7 days. At 7 dpi, lysates and organoids were harvested for downstream analysis. **(C).** RT-qPCR analysis of DIV35 organoids. The data are normalized to mock-infected cells and are presented as relative expression of EV-D68 to *GAPDH* with Mock set to 1. Data are presented as means ± SEM (n = four biological replicates). **(D).** Plaque-forming assay of supernatants from DIV35 organoids. Data are presented as means ± SEM (n = three biological replicates). ND = not detected; LOD = limit of detection **(E).** RT-qPCR analysis of RNA harvested of DIV85 organoids. The data are normalized to mock-infected cells and are presented as relative expression of EV-D68 to *GAPDH* with Mock set to 1. Data are presented as means ± SEM (n = three biological replicates). **(F).** Plaque-forming assay of supernatants harvested from DIV85 organoids. Data are presented as means ± SEM (n = three biological replicates). **(G).** Immunofluorescence images of *DIV35* forebrain organoids mock-infected or infected with either MO or Fermon viruses harvested at 7 dpi. Dashed boxes indicate the location of the zoom images. DAPI (blue) stains nuclei and VP1 (green, virus protein) indicates infected cells. **(H).** Immunofluorescence images of *DIV85* forebrain organoids mock-infected or infected with either MO or Fermon viruses harvested at 7 dpi. Dashed boxes indicate the location of the zoom images. DAPI (blue) stains nuclei and VP1 (green, virus protein) indicates infected cells. Scale bar - 100 µm. Statistical analysis performed with one-way ANOVA.

### Neuropathogenic and non-neuropathogenic EV-D68 strains rely on sialic acid for entry into forebrain organoids

Although an EV-D68 cellular receptor is still undefined, sialic acid or a sialylated glycoprotein has been proposed to be an entry receptor [18]. To determine whether differential levels of sialic acid expression could explain the divergent virus susceptibility between early and late organoids, we performed click chemistry. We found that both early and late forebrain organoids expressed similar levels of sialic acid **(Figure 3A)**. To demonstrate if EV-D68 entry into the forebrain organoids is dependent upon sialic acid, we treated organoids with neuraminidase, a sialic acid cleaving enzyme, for one hour prior to infection and then mock-infected or infected the organoids with either Fermon, MO, or MD-09/23229 (MD) viruses. MD is an EV-D68 strain that was isolated from a patient with respiratory illness in 2009, prior to the first reported AFM outbreaks in 2014, and represents a non-neuropathogenic strain in circulation independent of reported AFM cases [11, 19]. At 24 hpi, we harvested lysates and supernatants to measure virus entry. We found that in the early organoids, treatment with neuraminidase reduced viral copies **(Figure 3B)** of MO and MD strains. When we treated and infected late organoids, viral copies of both Fermon and MO were reduced **(Figure 3C)**. Further, only infectious viral titer of the MO strain was impacted by treatment with neuraminidase in either early or late organoids **(Figure 3D, 3E).** We also tested for the presence of an additional proposed EV-D68 receptor, intracellular adhesion molecule-5 (ICAM-5) [20]. We detected minimal ICAM-5 protein via immunofluorescence at either time point, and there was no statistical detectable difference in *ICAM-5* transcripts between the time points **(Supplemental Figure 3)**. Although forebrain organoids at both time points express sialic acid, the presence of sialic acid alone does not explain the strain differences observed.

**Figure 3.**
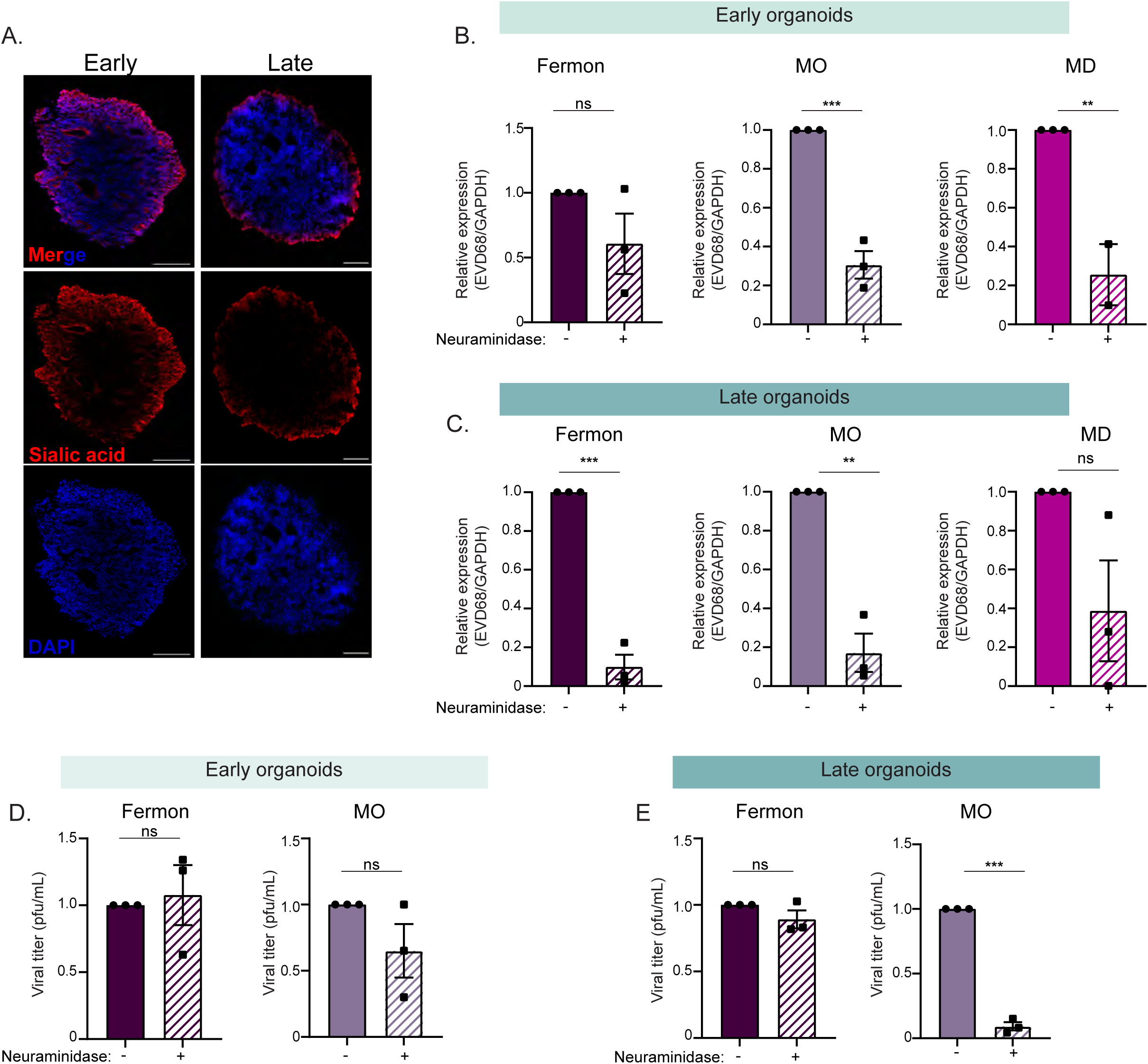
Enterovirus-D68 replication in forebrain organoids is partially dependent upon sialic acid expression. **(A).** Immunofluorescence images of DIV35 or DIV85 forebrain organoids for sialic acid protein expression using the Click-iT reaction. Scale bar - 200 µm **(B).** RT-qPCR analysis of lysates harvested from DIV35 forebrain organoids treated with neuraminidase for 1 hour and then infected with Fermon, MO, or MD viruses for 24 hours. **(C)** RT-qPCR analysis of lysates harvested from DIV85 forebrain organoids treated with neuraminidase for 1 hour and then infected with Fermon, MO, or MD viruses for 24 hours. The data are normalized to untreated organoids and are presented as relative expression of EV-D68 to *GAPDH* with untreated set to 1. Data are presented as means ± SEM (n = three biological replicates). **(D-E)** Plaque-forming assay of supernatants harvested at 1 dpi from forebrain organoids mock-infected or infected with either Fermon, MO, or MD at DIV35 **(D)** or DIV85 **(E)**. Data are presented as means ± SEM (n = three biological replicates). Statistical analysis was performed with unpaired student’s t-test.

### Early and late forebrain organoids contain distinct neural stem cell populations

Mammalian cortex development begins *in utero* as distinct laminations, consisting of six layers with marked neural cell types emerging throughout each layer [15]. Forebrain organoids recapitulate this anatomical lamination and cellular heterogeneity observed in the human brain, and thus serve as an ideal model to interrogate host responses in different neural contexts [13].

Early in forebrain organoid development, neural stem cells reside along the periphery of the organoid in a cellular niche called a neural rosette, analogous to the neural tube in the developing human cortex. As forebrain organoids mature, neural rosettes decrease, as developmentally mature neurons increase **(Figure 4A)**. To determine whether early and late forebrain organoids display differences in morphological features, namely the presence of these neural rosettes and mature neurons, we immunostained for SOX2 and MAP2, respectively. We found that early forebrain organoids display clusters of neural rosettes positive for SOX-2, a broad marker of neural stem cells **(Figure 4A)**. However, in late forebrain organoids, localization of SOX-2-positive cells shifts with a marked change in distinct pockets of accumulation. **(Figure 4A).** Both early and late forebrain organoids contain MAP2-positive cells, suggesting that differences between early and late development timepoints are largely centered around localization and accumulation of neural stem cells **(Figure 4A)**. Indeed, in our forebrain organoids, we did not detect any GFAP-positive astrocytes in our early organoids, in contrast to some visible GFAP-positive staining in late organoids **(Supplemental Figure 2)**, supporting what has previously been reported [17, 21, 22] Since we observed increased viral copies and infectious virus in the late organoids, but minimal GFAP-positive cells, immune signaling by astrocytes cannot account for the viral strain differences.

**Figure 4.**
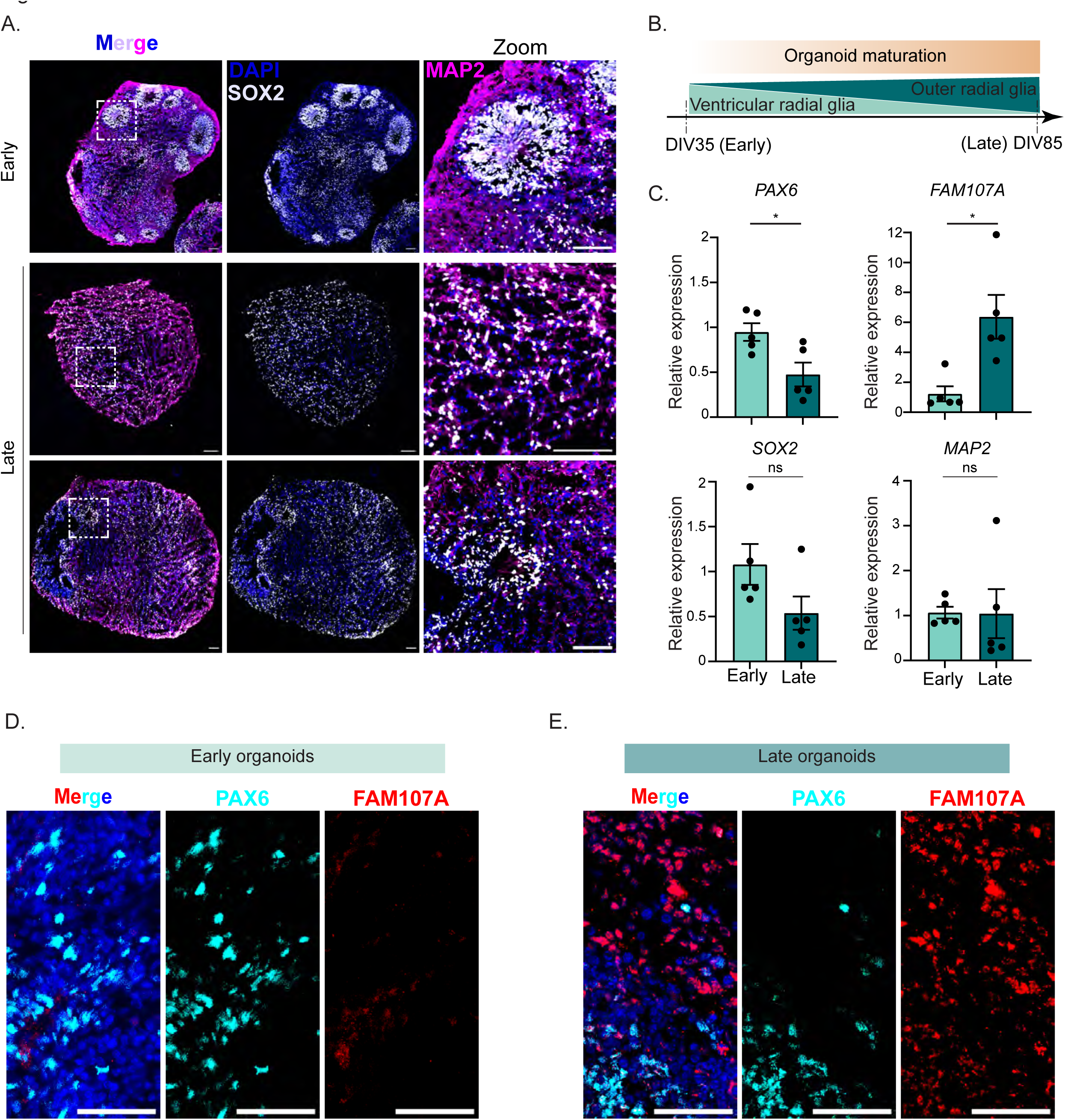
Early and late forebrain organoids display different morphology, protein expression, and protein localization. (**A).** Immunofluorescence images of early and late forebrain organoids. DAPI (blue) stains nuclei, MAP2 (magenta) stains maturing neurons, and SOX2 (grey) indicates neural stem cells. Scale bar - 50 µm **(B).** Schematic of cell type expression as organoids mature from DIV35 to DIV85 **(C).** RT-qPCR analysis of RNA harvested from uninfected forebrain organoids at DIV35 (early) and DIV85 (late). The data are normalized to the early samples and are presented as relative expression of the respective genes to *GAPDH*. Data are presented as means ± SEM (n = five biological replicates). Statistical analysis was performed with unpaired student’s t-test. **(D-E).** Immunofluorescence images of early **(D)** and late **(E)** uninfected forebrain organoids. DAPI (blue) stains nuclei, PAX6 (cyan) marks ventricular radia glia cells, and FAM107A (red) indicates outer radial glia cells. Scale bar – 50 µm.

The developing human cortex consists of several types of neural stem cells, named after the distinct cortical regions from which they arise. Important within the cortex are radial glia neural stem cells [23, 24]. Radial glia cells emanating from the ventricular zone are called ventricular radial glia cells (vRG) and those appearing in the outer subventricular zone are called outer radial glia (oRG) [25]. Interestingly, oRGs are human-specific, as rodent species do not contain them [26, 27]. oRGs emerge following vRGs, as the outer subventricular zone emerges later in development from the ventricular zone [28, 29]. As such, it has been shown that as forebrain organoids further develop *in vitro*, they begin to express more oRGs compared to vRGs and vice versa early in development ([30] **and Figure 4B)**. To confirm whether our early forebrain organoids contain increased vRGs compared to oRGs, we isolated RNA from uninfected early and late forebrain organoids and performed RT-qPCR for mRNA transcripts of the vRG marker, *PAX6*. We found that early organoids contained higher PAX6 transcripts compared to late organoids **(Figure 4C)**. Conversely, we found that late organoids express higher amounts of *FAM107A* **(Figure 4C)**. Consistent with our immunofluorescence images, we did not observe a significant difference in RNA expression of MAP2 or SOX2, although SOX2 trended towards increased RNA expression in the early organoids **(Figure 4C)**. Orthogonally, we performed immunofluorescence for PAX6 and FAM107A protein. We observed that early organoids express PAX6 protein in some neural rosettes **(Figure 4D)**, with minimal FAM107A protein present. Late forebrain organoids, however, expressed FAM107A protein radially emerging outward from some PAX6-positive neural rosettes **(Figure 4D)**. Taken together, these data implicate the presence of vRGs in early forebrain organoids, with oRGs appearing more in late forebrain organoids.

### Radial glia cells express high transcript levels of antiviral genes

The immune capacity of vRGs and oRGs has not been explored. Given that only the MO EV-D68 strain produced infectious titer in early forebrain organoids while infectious virus production from Fermon and MO was observed in late organoids, we hypothesized that early forebrain organoids are protected from EV-D68 infection by elevated levels of antiviral genes. Further, this primed antiviral gene state may be driven by radial glia cells present in the early forebrain organoids. We sought to evaluate the potential for antiviral priming in radial glia using a previously published single cell RNA sequencing dataset. Pollen and colleagues isolated cells from the ventricular and subventricular zones of human fetal tissue [25]. We re-clustered these data and identified/found four distinct clusters **(Figure 5A)**. Using gene lists marking distinct neural subtypes [25], we defined the clusters as radial glia, neurons, interneurons and ipc? (Figure 5B). Corroborating our analysis, the radial glia markers *PAX6* and *FAM107A* were among the top expressed in the radial glia cluster (**Figure 5B, arrows**). We next performed differential expression analysis to identify genes unique to the radial glial cluster compared to the other cell clusters. Interestingly, we found several immune genes that were upregulated in the radial glial cluster. Specifically, NF-κB-related genes were highly abundant, including *IL6ST*, *TNFRSF19*, *LITAF* **(Figure 5C)**. Further, we also identified interferon-stimulated genes predominantly present in the radial glia cluster, including *IFITM3* and *AXL* **(Figure 5D)**. These data suggest that radial glia cells are primed to be in an antiviral state by basally expressing key antiviral immune genes

**Figure 5.**
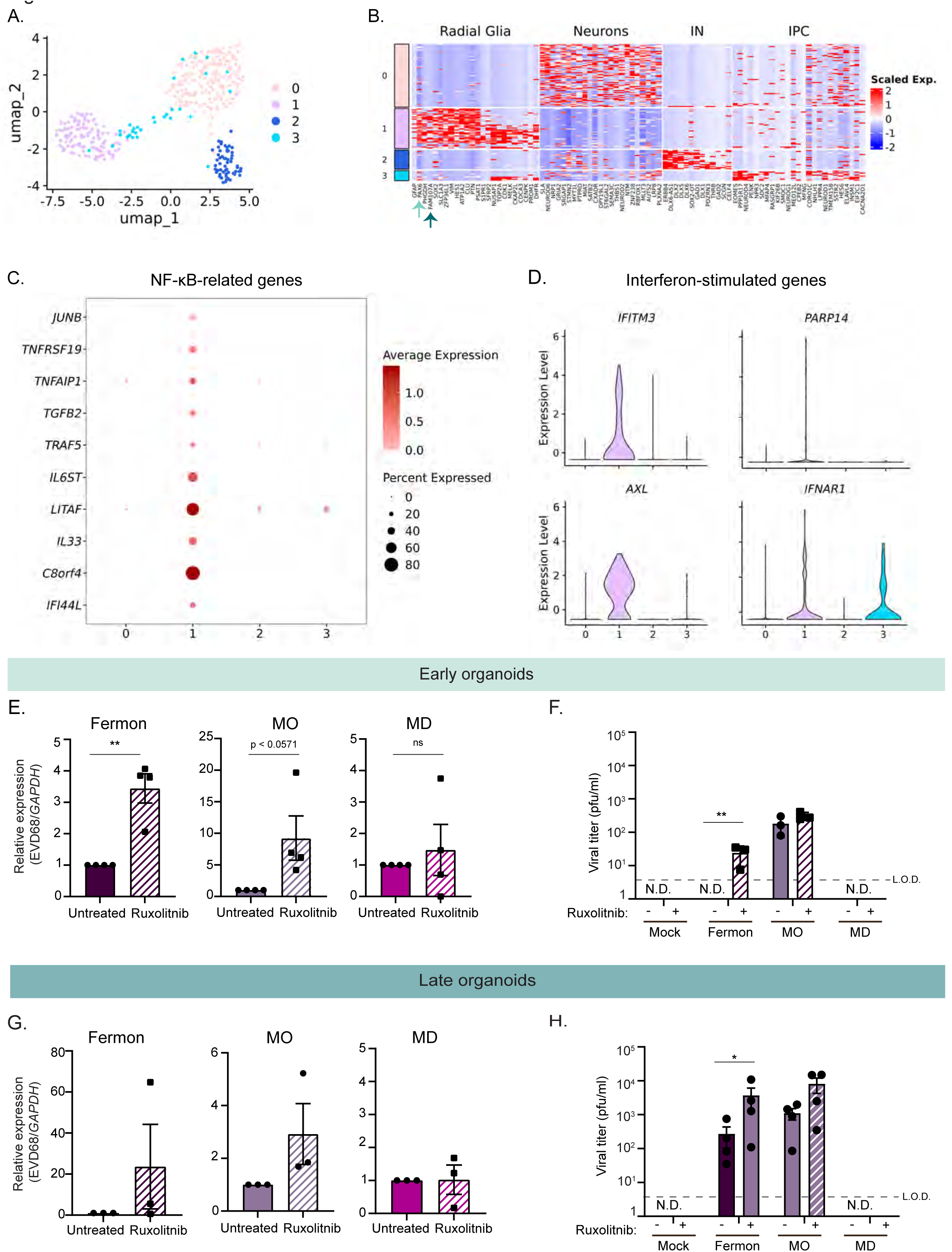
Differential interferon-stimulated gene expression during Enterovirus-D68 infection of early and late forebrain organoids. **(A)** Uniform manifold approximation and projection (UMAP) of previously published single cell transcriptomics data representative of 393 cells from ventricular and sub-ventricular zone human fetal brain samples [25]. **(B)** Heatmap displaying distinct neural cell populations. Arrows highlight radial glia specific genes. **(C-D)** Dot plot displaying NF-kB genes **(C)** and violin plots displaying interferon stimulated genes (ISGs) **(D)** identified using differential gene expression analysis of cluster 1 (radial glia) compared to other clusters (cutoff = log_2_ fold-change=1.9). **(E)** RT-qPCR analysis of lysates harvested at 7 dpi from DIV35 forebrain organoids untreated or treated with Ruxolitnib and simultaneously infected with Fermon, MO, or MD viruses. The data are normalized to untreated organoids and are presented as relative expression of EV-D68 to *GAPDH* with untreated set to 1. Data are presented as means ± SEM (n = four biological replicates). Statistical analysis performed with unpaired student’s t-test. **(F)** Plaque-forming assay of supernatants harvested at 7 dpi from forebrain organoids mock-infected or infected with either Fermon, MO, or MD at DIV35. Data are presented as means ± SEM (n = three biological replicates). Statistics performed with two-way ANOVA. N.D. = not detected **(G)** RT-qPCR analysis of lysates harvested at 7 dpi from DIV85 forebrain organoids untreated or treated with Ruxolitnib and simultaneously infected with Fermon, MO, or MD viruses. The data are normalized to untreated organoids and are presented as relative expression of EV-D68 to *GAPDH* with untreated set to 1. Data are presented as means ± SEM (n = three biological replicates). Statistical analysis performed with unpaired student’s t-test. **(H)** Plaque-forming assay of supernatants harvested at 7 dpi from forebrain organoids mock-infected or infected with either Fermon, MO, or MD at DIV85. Data are presented as means ± SEM (n = four biological replicates). Statistical analysis performed with two-way ANOVA. N.D. = not detected

### Immune activation limits EV-D68 infectious virus production in early and late forebrain organoids

Based on our identification of genes from multiple antiviral immune pathways present in radial glia cells and the previously established importance of the immune transcription factor STAT3 in neural stem cell maintenance and outer radial glia identity [25], we hypothesized that immune activation within forebrain organoids may limit EV-D68 infection. To this end, we utilized the JAK1/JAK2 inhibitor Ruxolitnib to broadly suppress immune activation. We simultaneously mock-infected or infected early forebrain organoids with either Fermon, MO, or MD viruses and treated with Ruxolitnib, at a concentration that has previously been used in other brain-derived organoids [31]. We found that JAK1/JAK2 inhibition resulted in increased virus RNA copies of Fermon and MO, but not that of MD **(Figure 5E).** We also found that infectious virus production increased with Ruxolitnib treatment, which is most evident during Fermon infection, in which infectious virus production is now detected and above the limit of detection **(Figure 5F)**. We performed the same infections and Ruxolitnib treatment in late forebrain organoids. Surprisingly, we found that viral copies in late organoids were not significantly increased during either Fermon, MO, or MD viruses **(Figure 5G).** Instead, we observed an increase in infectious virus production in the late organoids **(Figure 5H)**. These data emphasize the importance of evading neural stem cell immune activation, which may in turn, facilitate pathogenesis.

### Ventricular radial glia stem cells express antiviral proteins

Neural stem cells play an important role as the initiators of other neural cell lineages. As humans age, neural stem cell pools become depleted as a result of the emergence of mature neurons or glial cells. As such, protection of this limited cellular supply from viral infection, cellular death, and inflammation is paramount. One mechanism by which stem cells defend against pathogen insult is by upregulating various interferon-stimulated genes. Wu and colleagues found that neural stem cells basally express higher levels of IFITM family member proteins. We thus hypothesized that there may be additional ISGs that are basally upregulated in our early forebrain organoids. We extracted RNA from mock-infected early or late forebrain organoids and performed RT-qPCR for known antiviral ISGs [32]. We found that in addition to *IFITM1,* several other ISGs are upregulated in early organoids compared to late organoids **(Figure 6A)**. Interestingly, RNA expression of *ISG56* did not change in late organoids **(Figure 6A)**. We next hypothesized that our early forebrain organoids would express IFITM3 protein in PAX6-positive regions compared to our late forebrain organoids. We found IFITM3 protein expression in early organoids by both PAX6 expressing and not-expressing cells, while only PAX6-positive cells exhibited IFITM3 expression in late organoids **(Fig 6B)** In early organoids, we observed IFITM3 protein expression in PAX6-positive cells and also in PAX6-negative neural rosettes. In late organoids, IFITM3 protein expression was restricted to PAX6-positive cells **(Figure 6B)**. These data suggest that vRGs serve to defend against virus infection by upregulating IFITM3 expression, and that a decrease in PAX6-expressing cells and thus immune active cells, results in an increase in virus infectivity.

**Figure 6.**
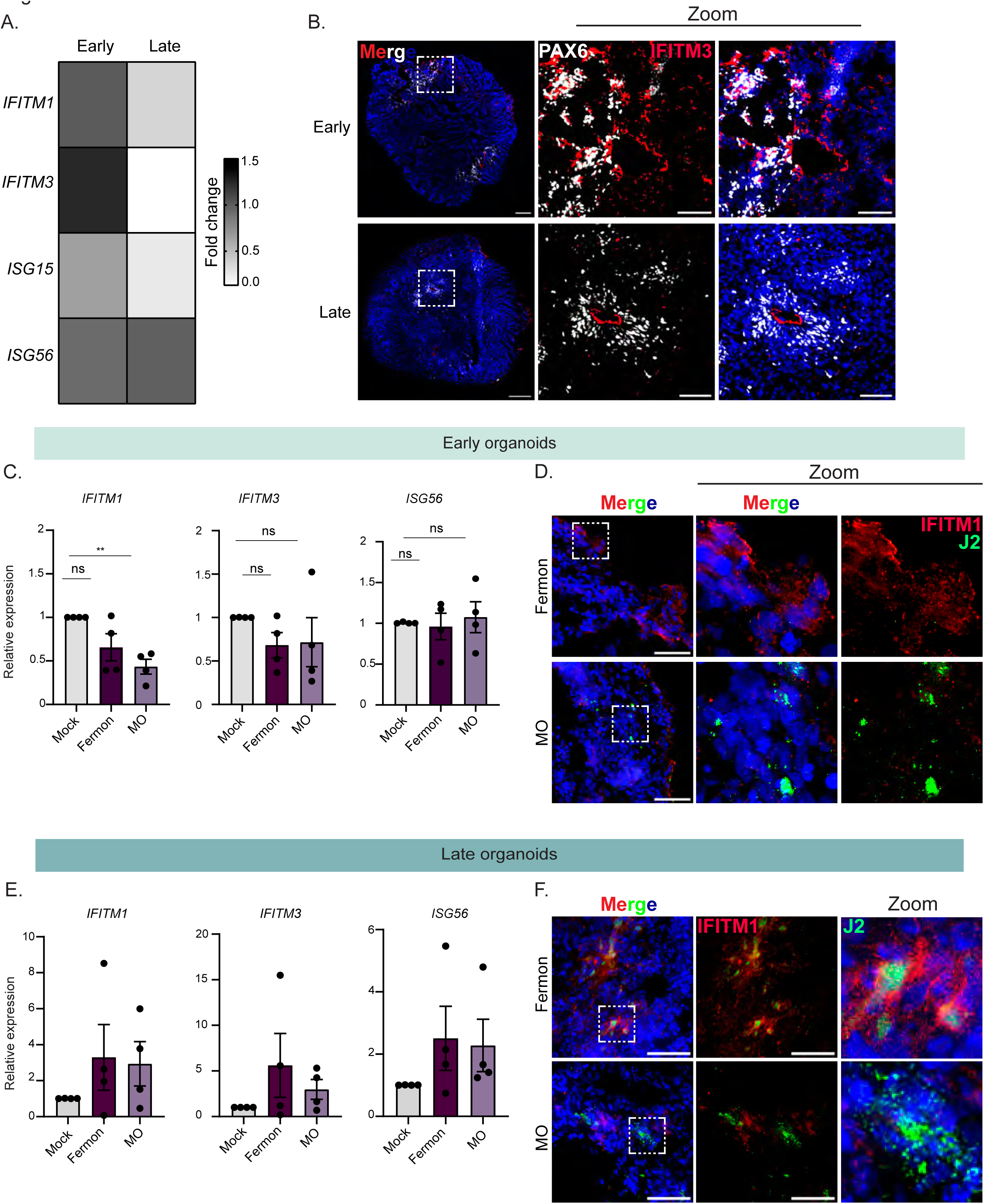
Distinct interferon signaling pathways are induced during Enterovirus-D68 infection of early and late forebrain organoids. **(A)** Heat map of fold change values from RT-qPCR of lysates from early and late forebrain organoids for several interferon-stimulated genes. **(B).** Immunofluorescence micrographs of early and late forebrain organoids. Organoids were stained for DAPI (nuclei, blue), PAX6 (grey, ventricular radial glia cells), and IFITM3 (red). Dashed boxes indicate the location of the zoom images. Scale bar (tiled image) – 200 µm; Scale bar (zoom image) – 25 µm. **(C).** RT-qPCR analysis of lysates harvested at 7 dpi from DIV35 forebrain organoids mock-infected or infected with Fermon, MO, or MD viruses. The data are normalized to mock-infected organoids and are presented as relative expression of the respective interferon-stimulated gene to *GAPDH* with mock-infected set to 1. Data are presented as means ± SEM (n = four biological replicates). **(D).** Immunofluorescence images of early organoids infected with Fermon or MO viruses at 7 dpi. Organoids were stained for DAPI (blue, nuclei), J2 (green, virus), and IFITM1 (red). Scale bar -50 µm. **(E).** RT-qPCR analysis of lysates harvested at 7 dpi from DIV85 forebrain organoids mock-infected or infected with Fermon, MO, or MD viruses. The data are normalized to mock-infected organoids and are presented as relative expression of the respective interferon-stimulated gene to *GAPDH* with mock-infected set to 1. Data are presented as means ± SEM (n = three biological replicates). **(F).** Immunofluorescence images of late organoids infected with Fermon or MO viruses at 7 dpi. Organoids were stained for DAPI (blue, nuclei), J2 (green, virus), and IFITM1 (red). Scale bar - 50 µm.

To determine whether virus infection within early forebrain organoids is dependent upon antagonizing this basally elevated interferon-stimulated gene signature, we measured RNA expression of several ISGs during infection with either MO or Fermon viruses. We found that MO reduced the induction and expression of *IFITM1*, compared to mock or Fermon-infected conditions **(Figure 6C)**. To identify IFITM1 antagonism on the protein level in single, infected cells, we performed immunofluorescence on MO or Fermon-infected early forebrain organoids staining for IFITM1 protein and an antibody against double-stranded RNA (J2). We found that in MO-infected cells, IFITM1 protein was not present, but we detected by increased IFITM1 protein expression in Fermon-infected organoids **(Figure 6D)**. Since Fermon does not productively infect early organoids, we did not detect Fermon-expressing cells, as demonstrated previously **(Figure 2G)**. This implies that at steady-state, neural stem cells express an antiviral program to protect the CNS, and for EV-D68 to establish a productive CNS infection, it must first overcome this upregulated ISG barrier.

## Discussion

In this study, we sought to identify cellular features mediating differential infectivity during EV-D68 infection within the CNS. We found that the canonical *in vitro* system used to study EV-D68 replication, SH-SY5Y cells, was permissive to both non-neuropathogenic and neuropathogenic EV-D68 strains, underscoring an alternative approach towards understanding EV-D68 infection within the CNS. Further, using forebrain organoids at two developmental time points, we identified that organoids at a later developmental stage were permissive to both neuropathogenic and non-neuropathogenic EV-D68 strains; however, only neuropathogenic EV-D68 strains were able to productively infect forebrain organoids at an early developmental time point. We identified that specific neural stem cells differentially expressed key innate immune signaling pathways that may dictate neuron-specific EV-D68 infection. Notably, the type of neural stem cell is important, as ventricular radial glia cells expressed antiviral immune proteins compared to outer radial glia cells, and EV-D68 regulation of this antiviral program may be critical in establishing neurological disease.

Until recently, the innate immune signaling capacity of neurons and neural stem cells has been woefully underappreciated [33]. Neurons have largely been considered immune signaling incompetent, with a role ascribed to simply responding to cues from microglia or astrocytes [34, 35]. Recent literature has suggested that neural stem cells can mount protective type I IFN signaling responses [33, 36, 37]. Interestingly, several immune genes have been implicated in maintenance of neural stem cells identity as well as brain development, including LIF and STAT3. However, a role in immune function and antiviral signaling capacity within neural stem cells has largely been overlooked. Our work expounds upon the existing literature by providing insight into immune pathways that are differentially expressed in neural stem cells compared to maturing neurons. Our study further illuminates the under-appreciated immune signaling capacity of neurons and how it interplays with susceptibility to virus infection.

The full cellular tropism of EV-D68 is largely unknown. Mouse model studies have demonstrated a preference for EV-D68 infection within spinal cord motor neurons. *In vitro* studies have also identified EV-D68 infection with cortical neurons and astrocytes, suggesting that EV-D68 could infect more than just spinal cord motor neurons [4, 6] To our knowledge, there is one documented fatal case of EV-D68-associated AFM in the United States, a 5 year old boy who was later found to have detectable EV-D68 viral RNA in the CSF [38]. Later, in a *New England Journal of Medicine* report, Vogt and colleagues obtained an autopsy sample from that patient and identified EV-D68 viral RNA and protein within the neurons of the anterior horn of the spinal cord [39]. Additionally, a recent study using human iPSC-derived spinal cord organoids found that only contemporary neuropathogenic EV-D68 strains, not Fermon, can infect these organoids at 14 days post-differentiation [40]. An outstanding question is whether Fermon is capable of infecting spinal cord organoids during later stages of development.

Viral determinants and host immune responses, likely together, contribute to the differences in neuropathogenesis of EV-D68. We show that viral factors promote infection within the CNS, as different non-AFM-associated EV-D68 strains (Fermon vs. MD) have vastly different CNS infection capacity. One potential reason why EV-D68 strain MD was not infectious in the forebrain organoids may be the temperature of the CNS (37°C vs 33°C). Though this assumption requires further investigation, it underscores the importance of evolutionary pressure in driving adaptation to the host and cellular environment. Given how relatively conserved the different EV-D68 strains are, it is interesting how much variation is observed in terms of replication kinetics and emergence of neuropathogenicity. Perhaps there are viral amino acid residues that mediate this variation or residues that persist despite the evolutionary pressure. Indeed, our amino acid alignment of the various EV-D68 strains (**Figure 1A)** reveal that the viral 3B protein sequence is fully conserved amongst many contemporary and historic EV-D68 strains. Additionally, mutations in the viral VP1 protein may also contribute to how EV-D68 establishes neuropathogenesis [41]. Future studies should examine the contributions of mutations in the viral life cycle, entry into the CNS, and immune evasion that allow viral persistence.

Though the host has several mechanisms by which it defends itself against virus infection, viruses have evolved ways to counteract such defenses. Our data suggest that limiting basal levels of immune genes may be one way viruses establish infection within nervous system tissue. Indeed, several viruses have been shown to preferentially infect neural stem cells, with Zika virus being one of the more well-characterized neurotropic viruses in recent years. Additionally, another study found that Japanese Encephalitis virus could replicate in less mature forebrain organoids but not in more mature forebrain organoids, even though there was a decrease in IFN induction in the later organoids [42]. This could suggest that the virus may encode viral proteins capable of immune antagonism. The same may hold true for EV-D68 as we know that EV-D68 can block several immune factors (IRF7 and TRIF) in non-CNS cells.

Overall, our work contributes to our understanding of differential immune regulation within neural stem cells, which can explain strain differences of neuropathogenic viruses.

### Limitations of study

This study largely focused on the capacity of the neurons and neural stem cells within the CNS to support and respond to EV-D68 infection. To probe neuron-intrinsic responses, we utilized a human induced pluripotent stem cell system. Though several studies have suggested that iPSC-generated neurons recapitulate human neurons, these cells likely differ in several aspects. Thus, future studies should examine immune responses and EV-D68 infection in primary neuronal cultures and postmortem patient samples.

## Acknowledgements

We would like to thank members of the Jurado laboratory and Ming laboratory for helpful discussion and manuscript review. We specifically would like to thank Ronni Kurzion, Angela Bongiovanni, and Emma LaNoce for technical support and guidance. We also acknowledge Dr. Scott Hensley and the Hensley laboratory for equipment usage. This work was supported by the University of Pennsylvania Office of the President (KAJ), the University of Pennsylvania Office of the Provost (CV and SGN), and the Burroughs Wellcome Fund (CV).

## Author Contributions

CV: Data curation, investigation, methodology, formal analysis, study conceptualization, experimental work, manuscript writing; SGN: Experimental work, formal analysis, manuscript review; CDB: Experimental work, manuscript review; SS: Experimental work; BH: Experimental work; G-LM: Methodology, study design, data interpretation, manuscript review; KAJ: Formal analysis, funding acquisition, supervision, study conceptualization, manuscript writing.

## STAR Methods

### Key Resources Table

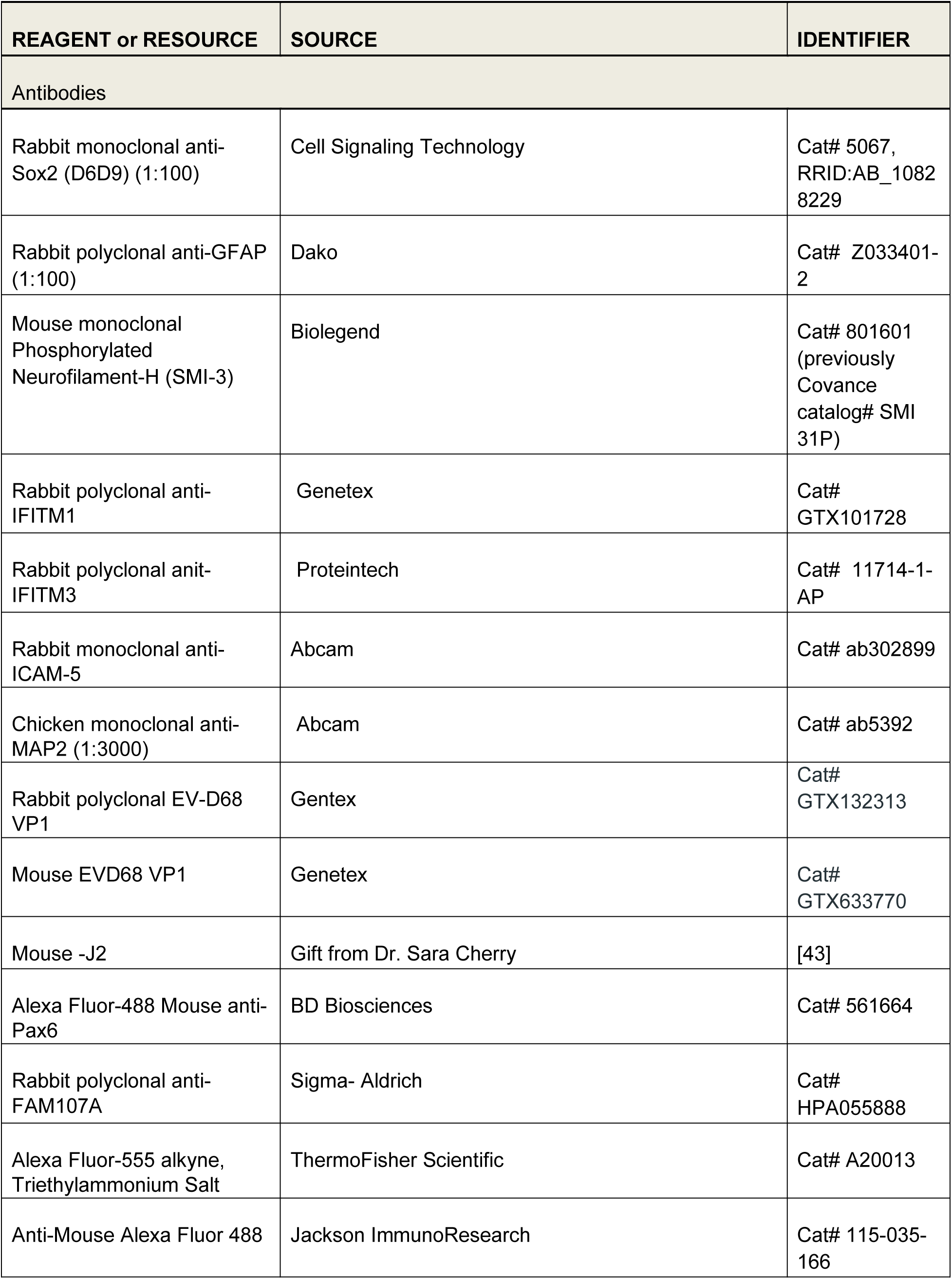
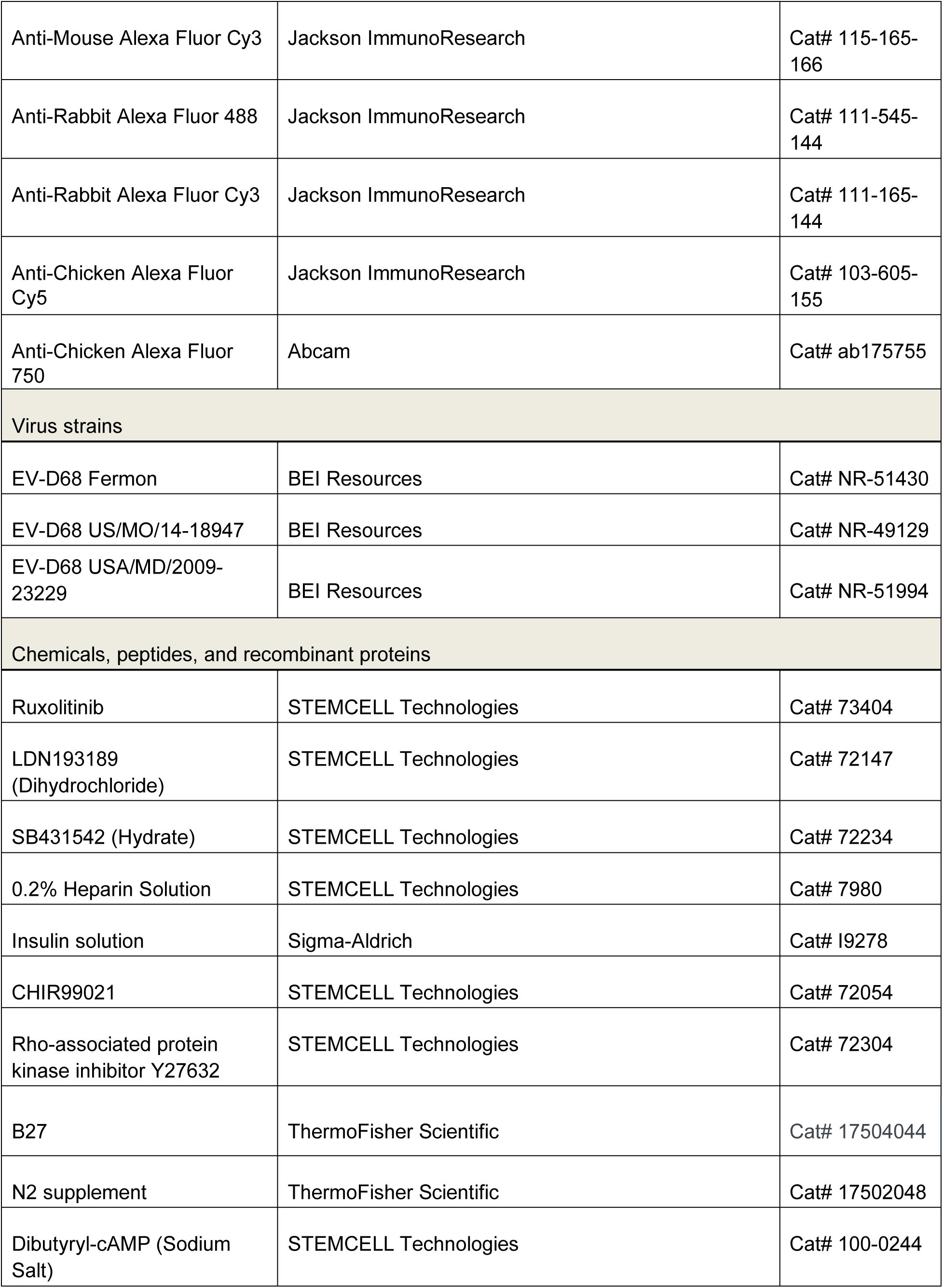
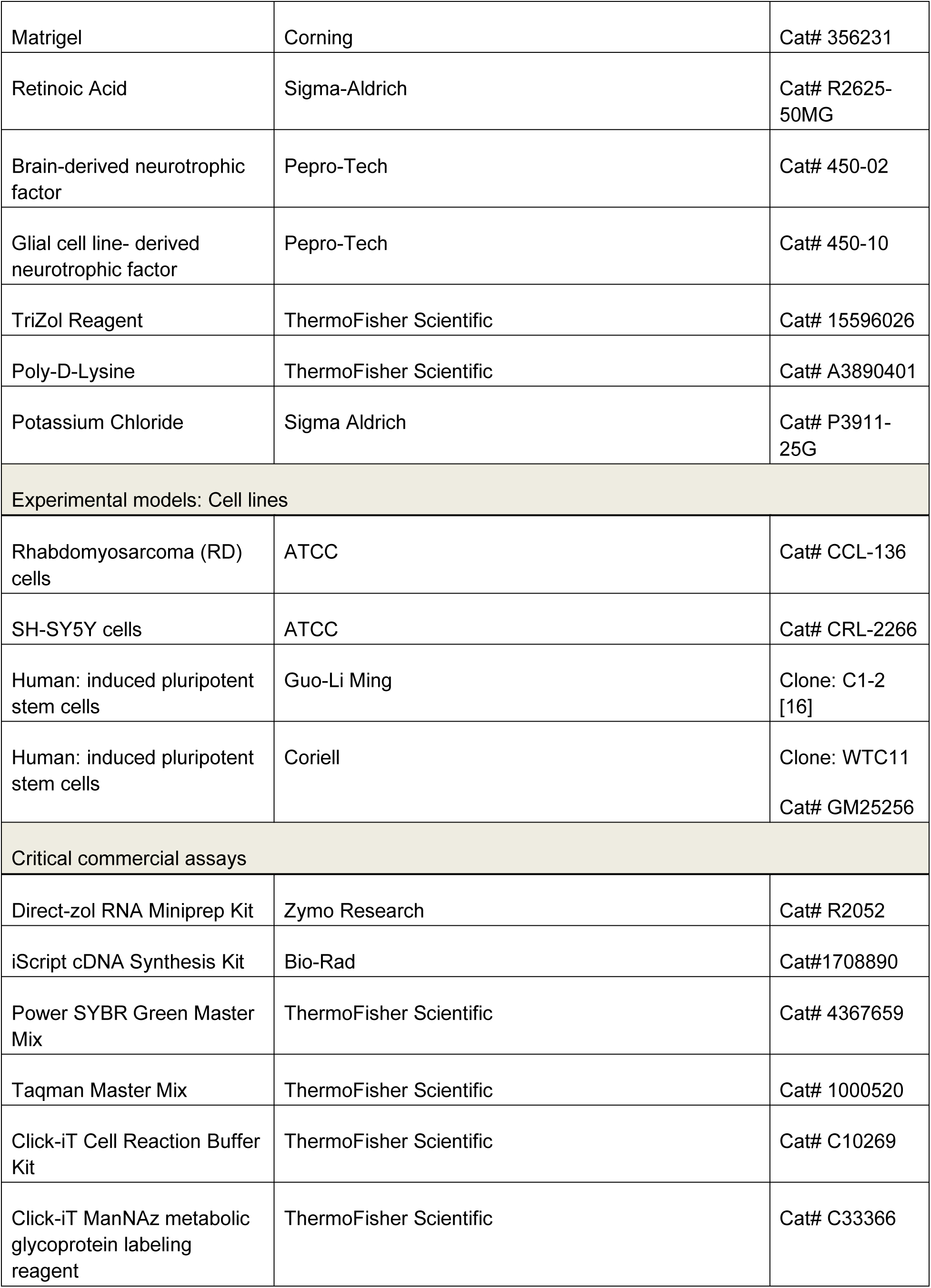
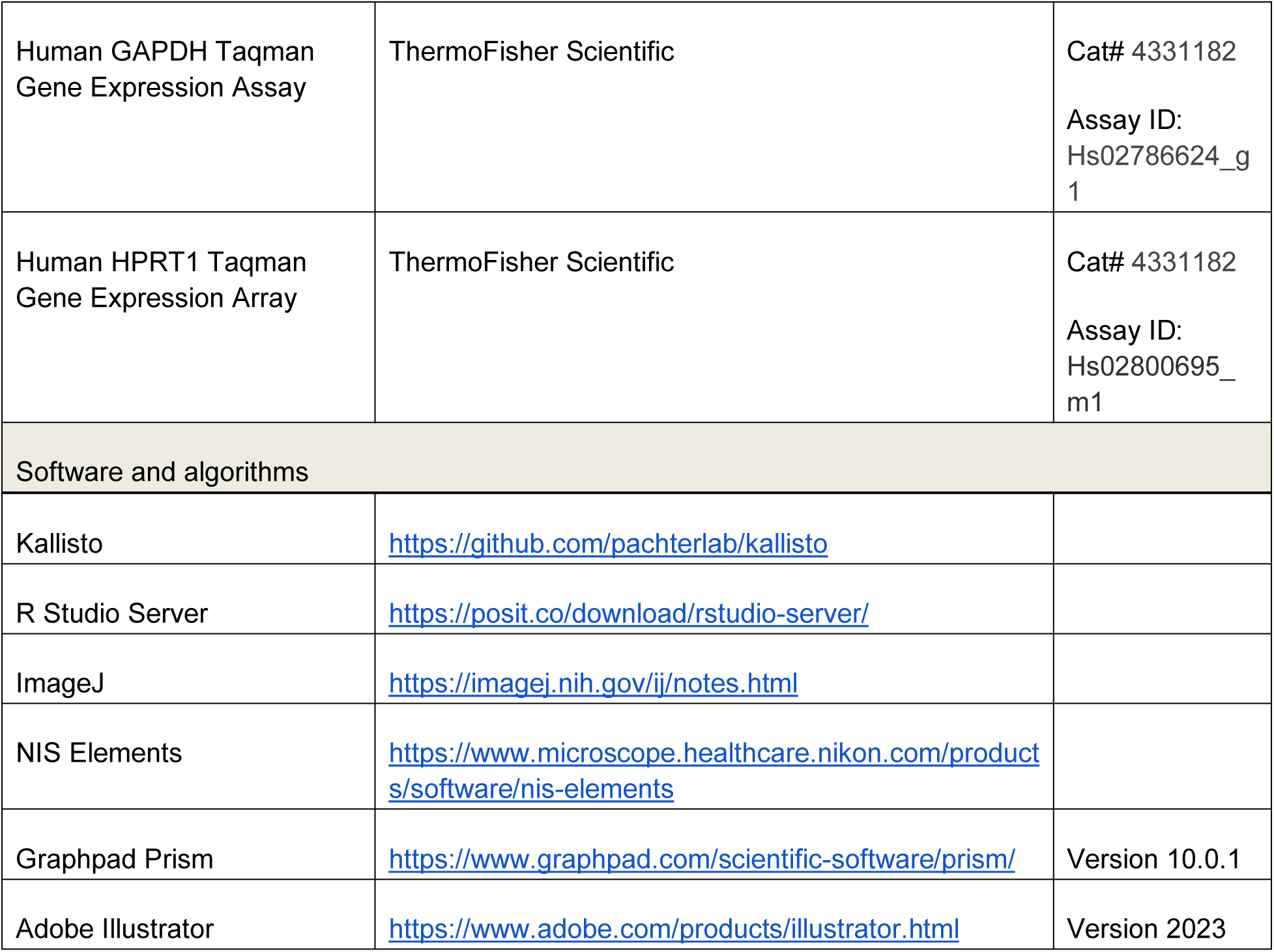

#### Lead contact

Further information and requests for reagents should be directed to and will be fulfilled by the lead contact, Dr. Kellie Jurado.

#### Cells and viruses

RD cells were grown in Dulbecco’s modification of eagle’s medium (DMEM; Gibco) supplemented with 10% fetal bovine serum (FBS; HyClone), 1% penicillin-streptomycin (Gibco), and 1% Gluta-Max (Gibco). Undifferentiated SH-SY5Y cells were maintained in DMEM/F-12/Glutamax medium (Gibco) supplemented with 10% fetal bovine serum (FBS; Hyclone). Enterovirus-D68 strains (Fermon, US/MO/14-18947, and MD/2009-23229) were purchased from BEI resources (catalog numbers NR-51430, NR-49129 and NR-51994, respectively). EV-D68 strains were propagated at the following temperatures: Fermon at 37°C, US/MO/14-18947 at 33°C, and MD/2009-23229 at 33°C. Human-induced pluripotent stem cell lines C1-2 and WTC11 (Coriell and [17]) were maintained in mTSeR media prior to cerebral organoid generation protocol. All cells were tested and found to be *Mycoplasma*-free at the Cell Center Services Facility (University of Pennsylvania).

#### SH-SY5Y cell differentiation

Differentiation of SH-SY5Y cells was performed as previously published for 18 days [10]. Briefly, undifferentiated SH-SY5Y cells were plated in six-well plates, followed by media changes with gradual depletion of FBS and addition of retinoic acid and a two-step cell splitting process. Initially, cells were maintained in differentiation media containing 2.5% heat-inactivated FBS and 10 µM retinoic acid. On day 7, cells were split 1:1 onto six-well plates, and the following day, differentiation media containing 1% heat-inactivated FBS and 10µM retinoic acid was added. On day 10, cells were split 1:1 onto poly-D-lysine-coated six-well plates. The following day, differentiation media containing 1X B27, 20 mM KCl, 50 ng/mL brain-derived neurotrophic factor, 2 mM dibutyryl cyclic AMP, and 10 µM retinoic acid was added. Media was changed every three days until day 18, at which point the cells were ready for experimental analysis.

#### Generation of forebrain organoids

Forebrain organoids were established following a previously established protocol [17]. Induced pluripotent stem cells were cultured with mTeSR Plus complete medium (STEMCELL Technologies) until ∼90% confluent. On day 0 of cerebral organoid generation, iPSCs were dissociated into a single cell suspension using ReLeSR (STEMCELL Technologies) and resuspended in mTeSR supplemented with Rho-associated protein kinase inhibitor Y27632 (ROCKi) at 10uM. non-cell culture treated 96-well U bottom plates were seeded with 200uL of cell suspension at a concentration of 2.5*10^5^ cells/mL (50,000 cells/well). Plates were incubated at 37°C and 5% CO_2_ for 2 days to form embryoid bodies (EB). On day 3 or organoid generation, EBs were resuspended and transferred into 6-well plates containing H1 medium (Basal Medium with F-12 nutrient (DMEM/F12), 20% Knockout Serum Replacement (KOSR), 1% glutamine supplement alternative (GlutaMAX), 1% MEM Non-essential amino acid solution (NEAA), 0.182% 2-beta-mercaptoethanol (BME), 1% penicillin/strep (Pen/Strep), 5uL of 1uM LDN, 25uL of 5uM SB, 0.2% heparin solution) supplemented with 10uM ROCKi. EBs were incubated at 37°C and 5% CO_2_ on a shaker at 120 revolutions per minute (rpm). On days 3-6, media was half-changed with H1 medium. On day 7, healthy EBs with round, smooth and bright edges were transferred into a Matrigel (ThermoFisher Scientific) and F2 medium solution (DMEM/F12, 1% N2 supplement, 1% GlutaMAX, 1% NEAA, 0.182% BME, 1% Pen/Strep, 5uL of both SB and CHIR at 1uM) and plated on ultra-low attachment 6-well plates. An EB/matrigel solution was incubated for 60 minutes at 37°C to solidify. F2 medium was added to the well and half-media changes were completed every other day until day 13 of organoid generation. On day 14, organoids (formerly EBs) were broken out of Matrigel and resuspended with H3 medium (47% DMEM/F12, 47% Neurobasal media, 1% N2 supplement, 2% B27 supplement, 1% GlutaMAX, 1% NEAA, 0.182% BME, 1% Pen/Strep and 12.5uL of 2.5ug/mL insulin) in 6-well plates. Plates were incubated at 37°C and 5% CO_2_ on a shaker at 120 rpm. Media changes were performed daily with H3 media until organoid harvest. For organoids that were maintained past DIV 70, media changes were performed with F4 medium (Neurobasal media, 2% B27 supplement, 1% GlutaMAX, 1% NEAA, 0.182% BME, 1% Pen/Strep, 5uL of 0.05mM cAMP, 5uL of 0.02mM ascorbic acid, 10uL of 20ng/mL BDNF and GDNF) and were sliced using a Leica VT 1200S vibratome between days 45-60 as previously described [15].

#### Plaque assay

Viral titer was determined using plaque assays. RD cells were plated on 6-well plates at a concentration that would allow for ∼80% confluency at the time of assay infection. Supernatants from infected cells or organoids, collected at the conclusion of respective experiments, were serially diluted and plated onto the RD cells. After virus adsorption for 1hr at 37°C, cells were overlaid with 1% SeaPlaque (Lonza), 1% SeaKem (Lonza) in MEM and 5% FBS. Plates were incubated at 37°C for 4 days, fixed with 10% neutral buffered formalin, agarose discs were removed, and plaques were stained with 0.5% crystal violet.

#### Immunofluorescence

Undifferentiated or differentiated SH-SY5Y cells were plated onto 6-well plates or 12-well plates and infected at varying multiplicities of infection the next day. Twenty-four hours post-infection, cells were fixed in 4% paraformaldehyde, permeabilized, and stained. For imaging of the forebrain organoids, organoids were fixed in 4% paraformaldehyde for 30 minutes with head-to-tail rotation, washed with 1X PBS, and incubated overnight in a 30% sucrose solution. Following sucrose incubation, the organoids were washed with 1X PBS, and mounted into cryomolds using TissueTek (Sakura Finetek). Organoids were sectioned into 10 µm sections using a Leica CM1950 or a Leica CM3050S cryostat, permeabilized, and stained. Images were acquired using a Nikon Ti2E scope. Image processing was conducted using ImageJ software.

#### Neuraminidase treatment

At either DIV35 or DIV85, organoids were incubated with neuraminidase [Sigma] at 6 units/mL for 1 hour. After 1 hour, organoids were washed with 1X PBS, and infected with EV-D68 strains or mock-infected. At 24hpi, organoids were washed with 1X PBS and harvested for downstream analysis.

#### Click-iT analysis

At DIV35 or 85, organoids were incubated with 25 µM Click-iT® ManNAz (tetraacetylated N-azidoacetyl-D-mannosamine) for 24 hours. Following incubation, organoids were washed with 1X PBS, fixed with 4% paraformaldehyde for 30 minutes - 1 hour, and incubated overnight in a 30% sucrose solution. Fixed organoids were placed in TissueTek cryomold and sectioned onto glass slides using a cryostat. The Click-iT analysis was performed on the glass slides according to the manufacturer’s protocol. Organoid sections were incubated in a solution of 1X Click-iT cell reaction buffer, CuSO_4_, Click-iT cell buffer additive, and 1 µM of 555-alkyne secondary antibody (ThermoFisher) for 30 minutes. Sections were washed with 1% BSA in PBS, incubated with 1X DAPI in PBS for 10 minutes, washed again with 1% BSA in PBS, and mounted with cover slips using Prolong Diamond Antifade Mountant (ThermoFisher Scientific). Image analysis on ImageJ was done by setting an organoid sample incubated without a label as the background reference and applying the LUTs settings to the organoids with the label.

#### RT-qPCR

RNA was extracted from cells or organoids using a Zymo Research Direct-zol RNA MiniPrep kit, and cDNA synthesis was performed on extracted RNA using iScript (Bio-Rad). The resulting cDNA was diluted 1:3 in ddH2O. RT-qPCR analysis was performed using either the Power SYBR green PCR master mix (Thermo Fisher) or Taqman Fast Advanced master mix (Thermo Fisher) on the QuantStudio 3 RT-PCR system. Primer sequences used are listed in **Table 1**.

**Table 1:**
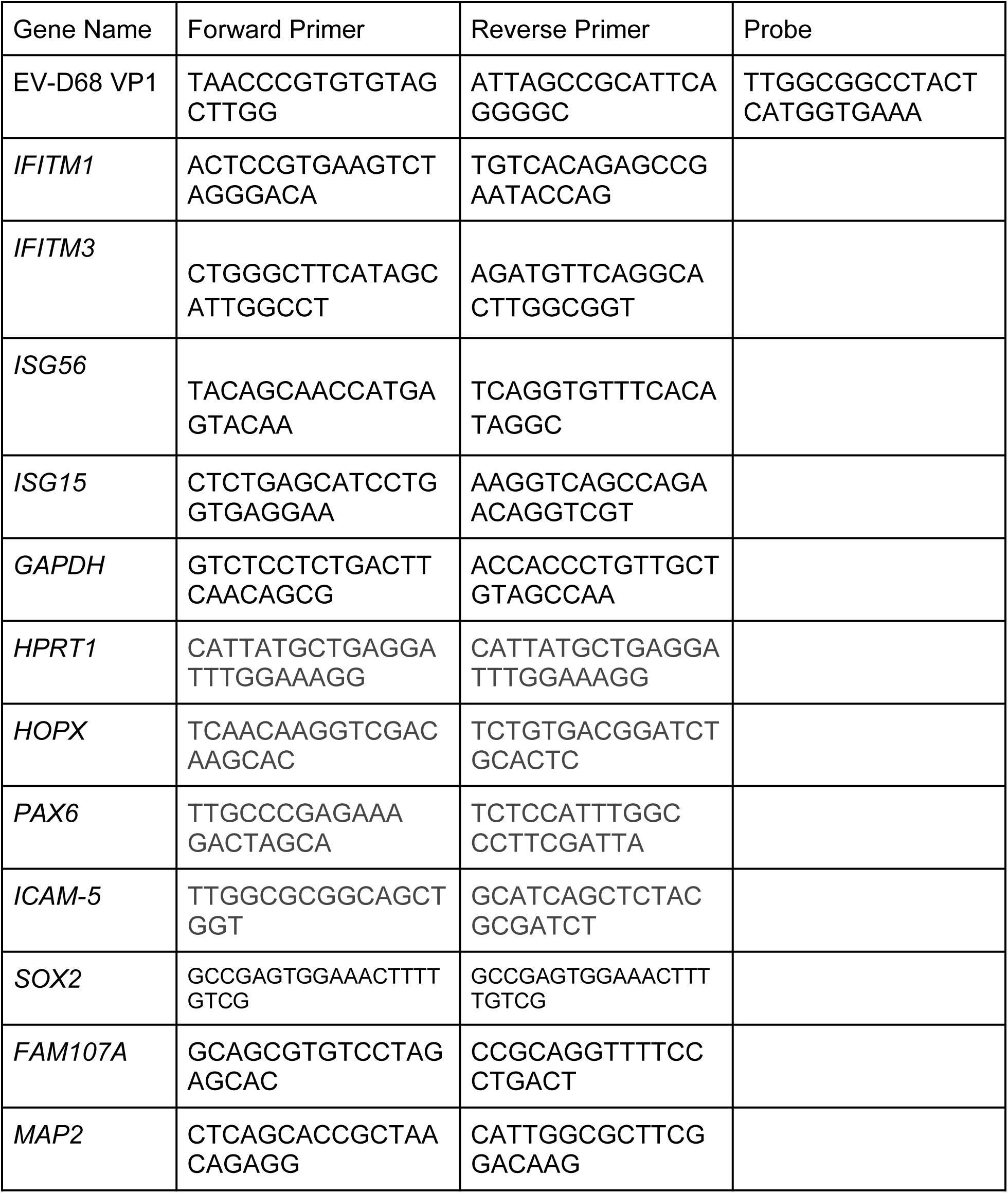
Primer sequences used in this study.

#### Single cell RNA sequencing analysis

Single cell transcriptomics data representative of cells isolated from the ventricular and sub-ventricular zone human fetal brain samples were obtained from a previously published dataset [25]. Specifically, we analyzed normalized gene expression values expressed as counts per million reads for all genes across 393 cells using R Studio Server. Using the Seurat package, we applied a principal component cut-off of 4 and a clustering resolution of 0.5 to identify 4 clusters. Using the previously defined cell identity markers, we generated a heatmap displaying each neural subtype. We next used Seurat’s FindAllMarkers to conduct differential expression analysis on radial glia and displayed NFkB-related and interferon stimulated genes using Seurat.

## Statistical analysis

All data were analyzed using the appropriate statistical test using GraphPad Prism software. One-way analysis of variance (ANOVA), two-way ANOVA, or unpaired student’s t-test were implemented for statistical analysis of the data using GraphPad Prism Software as indicated. Graphed values are presented as mean ± SEM (n = 3 or greater); *p ≤ 0.05, **p ≤ 0.01, and ***p ≤ 0.001.

---

**Supplemental Figure 1.**
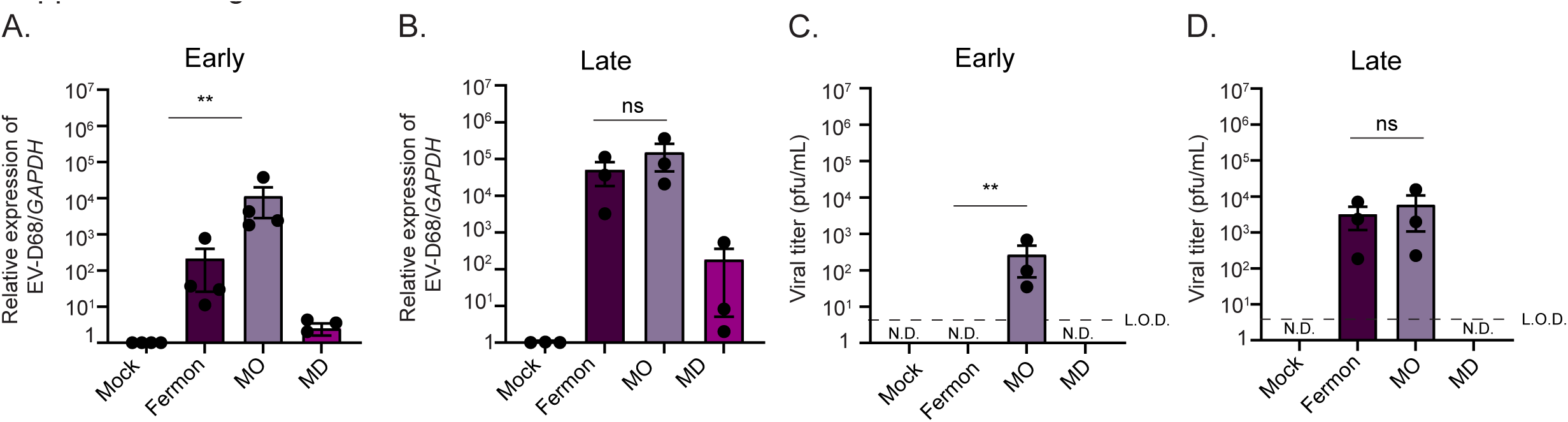
Forebrain organoids derived from a second human induced pluripotent stem cell line also support pathogenic Enterovirus-D68 infection. **(A)** RT-qPCR analysis of lysates harvested at 7 dpi from DIV35 forebrain organoids mock-infected or infected with Fermon, MO, or MD viruses. The data are normalized to mock-infected organoids and are presented as relative expression of EV-D68 to *GAPDH* with mock-infected set to 1. Data are presented as means ± SEM (n = three biological replicates). **(B)** Plaque-forming assay of supernatants harvested at 7 dpi from forebrain organoids mock-infected or infected with either Fermon, MO, or MD at DIV35 **(C)** RT-qPCR analysis of lysates harvested at 7 dpi from DIV85 forebrain organoids Fermon, MO, or MD viruses. The data are normalized to mock-infected organoids and are presented as relative expression of EV-D68 to *GAPDH* with mock-infected set to 1. Data are presented as means ± SEM (n = three biological replicates). N.D. = not detected **(D)** Plaque-forming assay of supernatants harvested at 7 dpi from forebrain organoids mock-infected or infected with either Fermon, MO, or MD at DIV85. Statistical analysis performed with one-way ANOVA. N.D. = not detected

**Supplemental Figure 2.**
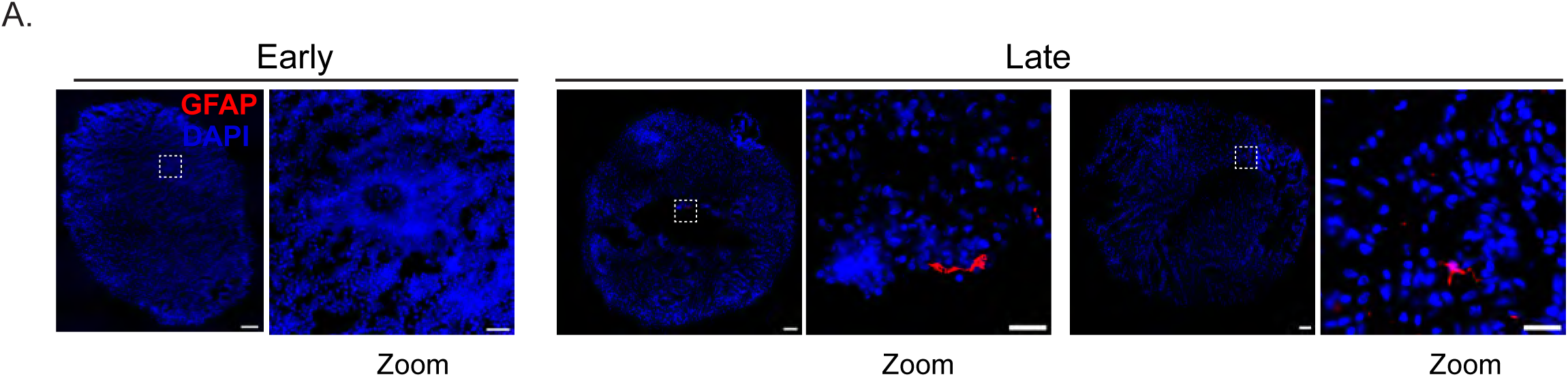
Late forebrain organoids contain few astrocytes. Immunofluorescence micrographs of early or late forebrain organoids. Organoids were stained for DAPI (nuclei, blue) and GFAP (red, astrocytes). Dashed boxes indicate the location of the zoom images. Scale bar (tiled image) - 200 µm; Scale bar (zoom image) - 25 µm.

**Supplemental Figure 3.**
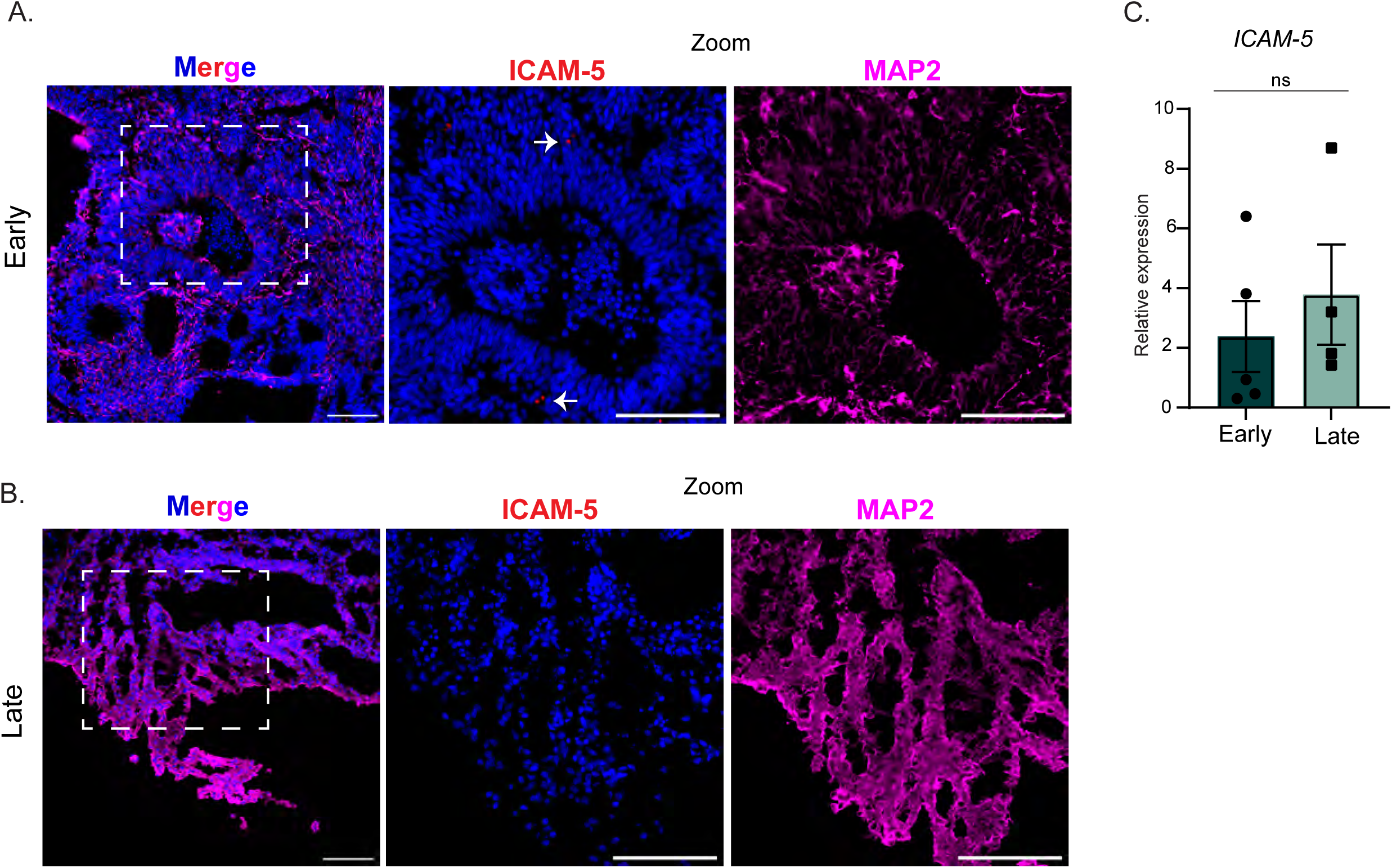
ICAM-5 is minimally expressed in forebrain organoids. Immunofluorescence micrographs of early **(A).** or late **(B).** forebrain organoids. Organoids were stained for DAPI (nuclei, blue), ICAM-5 (red), and MAP2 (magenta). Scale bar - 100 µm. **(C).** RT-qPCR analysis of RNA harvested from early or late forebrain organoids. The data are normalized to the early organoid sample and are presented as relative expression of *ICAM-5* to *HPRT1* with early set to 1. Data are presented as means ± SEM (n =five biological replicates). Statistical analysis performed with unpaired student’s t-test.

